# Airborne environmental DNA metabarcoding for the monitoring of terrestrial insects - a proof of concept

**DOI:** 10.1101/2021.07.26.453860

**Authors:** Fabian Roger, Hamid Ghanavi, Natalie Danielsson, Niklas Wahlberg, Jakob Löndahl, Lars B. Pettersson, Georg K.S. Andersson, Niklas Boke Olén, Yann Clough

## Abstract

Biodiversity is in decline due to human land use, exploitation, and climate change. To be able to counteract this alarming trend it is paramount to closely monitor biodiversity at global scales. Because this is practically impossible with traditional methods, the last decade has seen a strong push for solutions. In aquatic ecosystems the monitoring of species from environmental DNA (eDNA) has emerged as one of the most powerful tools at our disposal but in terrestrial ecosystems the power of eDNA for monitoring has so far been hampered by the local scale of the samples. In this study we report the first attempt to detect insects from airborne eDNA. We compare our results to two traditional insect monitoring projects (1) using light trapping for moth monitoring and (2) transect counts for the monitoring of butterflies and wild bees. While we failed to detect many of the same species monitored with the traditional methods, airborne eDNA metabarcoding revealed DNA from from six classes of Arthropods, and twelve order of Insects - including representatives from all of the four largest orders: Diptera (flies), Lepidoptera (butterflies and moths), Coleoptera (beetles) and Hymenoptera (bees, wasps and ants). We also recovered DNA from nine species of vertebrates, including frogs, birds and mammals as well as from 12 other phyla. We suggest that airborne eDNA has the potential to become a powerful tool for terrestrial biodiversity monitoring, with many impactful applications including the monitoring of pests, invasive or endangered species or disease vectors.

## 1 Introduction

Global biodiversity is declining at an unprecedented rate and more species are threatened with extinction than ever before (Brondizio et al. 2019). One of the hardest hit groups are terrestrial insects and a series of recent studies (Hallmann et al. 2017; Sánchez-Bayo and Wyckhuys 2019) reported such massive declines in insect biomass that the New York Times magazine on 27^th^ of November 2018 declared that “The Insect Apocalypse is Here” (Brooke 2018).

The most comprehensive assessment to date (van Klink et al. 2020) nuances earlier more drastic findings, but still estimates a sustained decline in terrestrial insect abundance of 9% per decade. The main message however lies in the uncertainty of these estimates themselves, which are inherent to the fact that for the vast majority of insect species the data simply do not exist. Globally, only ∼9000 insect species are listed on The Red List, representing ∼0.2% of an estimated 5.5 million species worldwide (Stork 2018) and even here one-third are categorized as data deficient (IUCN 2019). Despite their importance, data on the occurrence of insects are scarce for all but a few well-characterized taxa, and only a single systematic insect monitoring programme exists at European level - the European Butterfly Monitoring Scheme (Van Swaay et al. 2019). This lack of data impedes efforts to accurately quantify and follow the state of insect populations, a critical prerequisite to studying the consequences of anthropogenic pressures such as climate and land-use change. However, given the immense and often cryptic diversity of insects and the time and taxonomic skills required for a certain identification, traditional monitoring methods cannot fill this gap. New, cost-effective and scalable monitoring methods are needed.

In recent years, DNA-based methods have enabled significant progress in monitoring biodiversity of natural communities. DNA metabarcoding has been successfully employed for large bulk samples of arthropods (Yu et al. 2012; Ji et al. 2013) and shown to be faster and taxonomically more comprehensive than traditional morphometric identification. When three high-quality standard datasets which contained > 55000 arthropod and bird specimens were identified to species level by both metabarcoding and morphometric identification (the latter took 2505 work-hours) the metabarcoding approach proved superior (Brehm et al. 2016). However, studying insects through the metabarcoding of bulk samples, requires the species to be collected first and most trapping methods are only suitable for certain taxonomic or functional groups (Matos-Maraví et al. 2019). In addition, the method always necessitates the killing of all captured insects.

In contrast, the metabarcoding of extra-organismal environmental DNA (eDNA) is non-invasive and has revolutionized the detection of biodiversity in aquatic ecosystems (Deiner et al. 2017; Ruppert, Kline, and Rahman 2019). Here, species are detected from the traces they leave behind, including shedded cells, tissue fragments, body excretions or gametes (Taberlet et al. 2018). The technique has been proven very powerful for the detection of fish and amphibians but also for reptiles and aquatic insects (Deiner et al. 2017; Thalinger et al. 2021; Rees, Maddison, et al. 2014). A single-species assay for the detection of the great crested newt is the first eDNA based method approved by regulators (Rees, Bishop, et al. 2014) and eDNA surveys are successfully employed to track the progression of aquatic invasive species (Jerde et al. 2011; Larson et al. 2020; Mauvisseau 2019).

In terrestrial ecosystems, the deployment of eDNA methods for biodiversity surveillance has proven more difficult and the biggest challenge that terrestrial applications of eDNA face is the local and non-representative nature of the samples. For insects, flower swabs have been sequenced to study plant-pollinator interactions (Thomsen and Sigsgaard 2019), metabarcoding of cow dung has been used to study dung-associated invertebrates (Sigsgaard et al. 2021), Honey has been studied to detect sap-sucking insects (Utzeri et al. 2018) and Lin et al. (2020) analysed top-soil samples from across California and analysed it for arthropod DNA (among other groups). However, in contrast to water which carries the DNA from all organisms living in the aqueous matrix, these examples represent an interaction between a small number of species and/or have a local geographic signal. While top-soil can potentially represent a time-integrated sample of the diversity above it, it is very challenging to collect samples that are large enough to be representative for any given location and DNA extraction from large amounts of soil imposes additional difficulties (Taberlet et al. 2018). This makes the aforementioned approaches unsuitable for the biomonitoring of diverse groups of organisms or for surveying large geographic areas. Valentin et al. (2020) have proposed to overcome these challenges by spraying water on leaf surfaces to collect an aggregated eDNA sample from larger areas or to roll stems and twigs with cotton to aggregate eDNA in this way, but whether these techniques are suitable for collecting representative samples from the larger insect community remains to be shown.

### 1.1 Sampling eDNA from the air

Air, as water, carries a multitude of microscopic particles from the inorganic and organic environments, so called aerosols. Aerosols of biological origin are called bioaerosols and can be classified into three main fractions (V. Després et al. 2012; Šantl-Temkiv et al. 2020). (1) living organisms such as viruses, bacteria, microalgae and unicellular fungi (2) propagules (pollen or spores) released into the air by plants and fungi and (3) excretions and cell or tissue fragments form microbial life, animals or plants, on their own or bound to non-biological particles. The study of the first fraction (airborne microorganisms) is an established research field (aerobiology, Womack, Bohannan, and Green (2010); Smith, Griffin, and Jaffe (2011)) and metabarcoding of aerosol samples is an established practice (Després et al. 2007; Després et al. 2012; Behzad, Gojobori, and Mineta 2015). While most research has focused on bacteria, microalgae (Tesson et al. 2016) and fungi have also been explored (Fröhlich-Nowoisky et al. 2012). Plant pollen is intensively studied for its allergenic potential and public health implications and metabarcoding is starting to be used for its identification (Kraaijeveld et al. 2014; Brennan et al. 2019; Rowney et al. 2021). Fungal spores were targeted for biodiversity monitoring of the regional fungal community (Abrego et al. 2018; 2020) and are currently being evaluated worldwide in the Global Spore Sampling Project (Ovaskainen et al. 2020).

The third fraction of bioaerosols has the greatest potential to harbor eDNA that is representative of animal diversity at large, but as far as we can tell this is virtually unexplored. Only a single study, published this year, showed that eDNA from a vertebrate species could be collected from the air (Clare et al. 2021). While an interesting proof of concept, it was performed on rather artificial conditions, inside an animal housing room with high densities of the target species. The only studies to explicitly sample airborne eDNA other than pollen or spores in an outdoor environment for the detection of macroorganisms are Johnson, Cox, and Barnes (2019) and Johnson et al. (2021) who successfully detected the insect pollinated shrub honey mesquite outside of its flowering season. To the best of our knowledge no targeted detection of airborne insect DNA has been attempted to date.

Yet, auxiliary evidence exists: An electron microscopy study of aerosols collected on the city outskirt of Munich (Wittmaack et al. 2005), observed a variety of insect scales and the authors note that “amongst the various bioaerosols observed in this study, scales of insects are prominent in terms of size”, thus confirming the presence of insect body parts in air. It is noteworthy that the samples represented rather small volumes of air (14 L*mm^-2^ of filter) and that insect scales were observed more frequently than bacteria in this study. In one of the only studies targeting eDNA to study insect communities, Madden et al. (2016) analysed dust samples collected indoors, by citizen scientists in 700 households across the USA. The authors detected 600 genera of arthropods, belonging to 28 orders. Importantly, all detected eDNA in that study is likely to have been airborne before settling on the collection site (door rims). The most direct evidence however comes from two metagenomic studies targeting aerosols directly: Be et al. (2015) characterized the aerosols captured on air filters (collected by the biological agent surveillance program BioWatch - U.S. Department of Homeland Security) and found sequences mapping to invertebrates, including insects, were the third most abundant fraction after bacteria and plants, peaking in summer. Likewise, Yooseph et al. (2013) collected aerosols from indoor and outdoor environments which they analysed using shotgun metagenomic sequencing. The authors report the detection of 711 insect genera, belonging to 22 orders, as well as non-flying arthropods such as 45 genera of spiders (order Aranea).

In this study, we explored the potential of metabarcoding of airborne eDNA to monitor the diversity of terrestrial insects, investigating three key questions: (1) can insects be detected from airborne eDNA (2) are the results comparable to those of traditional sampling, and (3) is airborne eDNA representative of the local species community allowing to distinguish between different sampling locations and landscapes in terms of species composition and diversity. To this aim we collected aerosols at three sites, two of these were carried out in parallel to two ongoing traditional insect sampling schemes: (1) a moth monitoring survey and (2) a pollinator survey. To the best of our knowledge this is the first attempt to detect eDNA from insects in the air and the first attempt to detect airborne eDNA from any animal species in a natural environment.

## 2 Material and Methods

### 2.1 Aerosol sampling method

We collected airborne eDNA using the biological air sampler Coriolis (μ) (Bertin Instruments, Montigny-le-Bretonneux, France). The sampler is a liquid cyclone sampler, which collectes aerosols by swirling the air in a conical tube filled with a collection liquid. Due to their inertia, aerosols are transferred into the liquid by impaction while the air escapes upwards from the centre of the swirl. The sampling duration was 10 minutes at a flow rate of 300 l*min^-1^ equaling a total sampling volume of 3000 l or 3 m^3^ of air per sample. To further increase the amount of sampled air, we collected and pooled two to three consecutive samples, resulting in a total collection volume of 6-9 m^3^ air. To prevent small insects from getting sucked into the sampler, we placed a commercial butterfly net (Naturbokhandeln, Mörbylånga, Sweden) over the mouthpiece of the air sampler. We used 15 ml of Milli-Q water as collection liquid, which we transferred into a 50 ml Falcon tube directly after sampling. The samples were put on ice immediately and frozen at -20 °C the same day. To prevent cross-contamination, a new sterile collection cone was used for each sample and all parts in contact with the air during collection (the insect net, the mouthpiece of the coriolis sampler and the metal tube connecting the suction pump with the mouthpiece), were soaked in a 5% chlorine solution for a minimum of 10 min between each sampling, after which we thoroughly cleaned all pieces with deionized water and rinsed them with Milli-Q water before re-using. We wore nitrile gloves at all times and changed gloves between all samples.

### 2.2 Air sampling and traditional insect surveys

We collected aerosol samples at three locations in southern Sweden, two of which in parallel to two insect surveys, a Moth monitoring scheme and a Pollinator survey (called “Moth site” and “Pollinator site” henceforth). The Moth monitoring scheme was part of a systematic monitoring of Swedish moth species that has been ongoing for more than a decade (Franzén et al. 2020). The project monitors moths using an automatic light trap (Robinson type) with a 250 W mercury vapour lamp (Leinonen et al. 1998). The trap is located at approximately eight meters above ground level, on the rooftop on the western side of the Ecology Building at Lund University, Sweden (Lund: Sölvegatan 37, 55°42’51.21’’ N, 13°12’26.27’ E). The trap was lit during night and all macro moth species were determined to species level by taxonomic experts. In total, we collected ten aerosol samples in parallel to the moth trapping, on ten consecutive week-days between August 27^th^ and September 9^th^, 2019. To avoid sampling DNA carrying particles originating from the moth trap itself, the air sampling was not performed next to the trap but on a balcony on the south side of the building, at the same height. In addition, the sampling was conducted early mornings, before the light trap was opened and the moths released. We pooled three consecutive samples at each sampling location, yielding a sampling volume of 9 m^3^ of air.

The pollinator survey was part of a separate research project, where wild bees and butterflies were inventoried in different open habitats in a forest dominated landscape in Småland (southern Sweden). The survey inventoried pollinators on different types of land uses (habitats), nested in landscapes - where each landscape was defined as a patch of habitats associated with one or several farmsteds. The survey used a combination of line transects, with five transects per site, and free counts. The transects were distributed approximately evenly along the edge of the habitat and directed towards the centre. Each transect was 50 meters long and was walked along for 10 minutes, from the edge out in the habitat. All wild bee and butterfly species were identified to species level. Additionally, 10 minutes of free counts were done in areas considered to be of better quality within each habitat. The surveys were conducted between 9:00 a.m. to 6:00 p.m. on days with favourable conditions for bumblebees and butterflies, i.e. days without rain, temperatures above 17 °C and wind speeds not above 4 on a Beaufort scale. We visited three of the surveyed landscapes on August 5^th^ 2019 and collected aerosol samples on five sites in three different habitats in each landscape: two semi-natural pastures, three ley fields and one clear-cut that has been cut within the last 10 years. At each site we collected and pooled two aerosol samples, resulting in a sample volume of 6 m^3^ of air per sample and a total of 15 samples.

The third sampling location was a popular local recreational park-like area comprising beech-dominated forest and open areas (Bokskogen, near the city of Malmö). Here we collected an additional five samples on September 22^nd^, located within ∼1km of each other. We refer to the site as “Beech forest”.

As negative control, we collected two clean samples (3 m^3^) in a particle free hood at the Lund University Aerosol Laboratory (Department of Ergonomics and Aerosol Technology, LTH, Sweden). The samples were pooled and carried along through sequencing.

A map of all sampling locations is shown in Figure 1.

**Figure 1.**
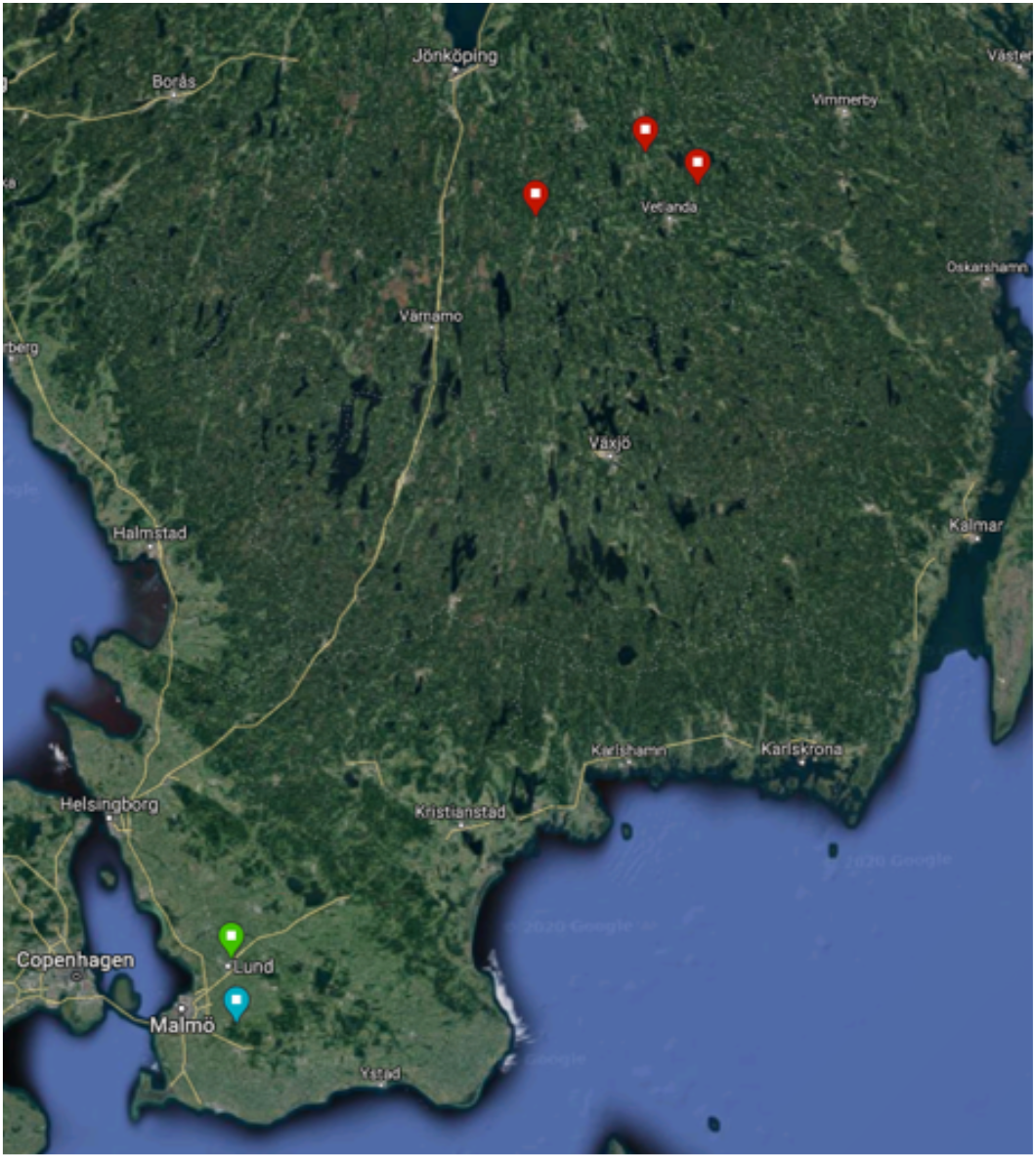
A map showing the three sampling locations in southern Sweden. The red marks designate the sampling location in Småland, the “Pollinator site”. Each red mark marks one sampling site from that scheme and five samples were collected at each of the three sites (all on the same day). The green mark marks the location of the “Moth site” (on the roof of the Ecology building from Lund University in the city of Lund). Here ten samples have been taken on ten consecutive week-days. The blue mark designates the third sampling location in a popular recreational beech forest area near the city of Malmö (“Beech forest”). Here, another five samples have been taken a few hundred meters apart, all on the same day.

### 2.3 DNA extraction, amplification and sequencing

The samples were thawed at 50°C and filtered manually onto a Millex-GV syringe filter with 0.22 µm pore-size and 25 mm in diameter hydrophilic polyvinylidene difluoride (PVDF) membrane, using a sterile 50 ml syringe. The filter was subsequently opened using plumbing pliers and the membrane was transferred with tweezers into a 1.5 ml collection tube. Filtering and DNA extraction was performed in a dedicated laboratory, free from post-PCR products. The working area was cleaned with a 5% chlorine solution before the start of the work and the pliers and tweezers were soaked in chlorine and rinsed with deionized water between each sample. We wore nitrile gloves at all times and changed gloves between each sample.

We extracted DNA with the Marcherey-nagel NucleoSpin Tissue kit (GmbH & Co. KG, Düren, Germany) following the manufacturer’s instructions for human or animal tissue and cultured cells. We eluted the samples twice (using 50 μl of BE Buffer each time) into the same collection tubes. The incubation time for each elution was five minutes, at room temperature.

We used two different metabarcoding primer pairs: BF3 **5’-CCHGAYATRGCHTTYCCHCG-3’** and BR2 **5’-TCDGGRTGNCCRAARAAYCA-3’** (Elbrecht and Leese 2017; Elbrecht et al. 2019), targeting the mitochondrial COI-gene and Chiar16SF **5’-TARTYCAACATCGRGGTC-3’** and Chiar16SR **5’-CYGTRCDAAGGTAGCATA-3’**(Marquina, Andersson, and Ronquist 2019), targeting the mitochondrial 16S-gene. Both primers have been developed for the metabarcoding of terrestrial insects. Optimal annealing temperature for both primer pairs was determined using a gradient PCR, amplifying a known lepidopteran DNA extraction, which suggested an optimal annealing temperature of 46° C for both COI and 16S primer pairs. The COI gene is the most common gene used for metabarcoding in the animal kingdom and is accompanied by a very comprehensive reference database, the BOLD database (Ratnasingham and Hebert 2007). However, there is evidence that even the best-performing primers fail to amplify many species of Hymenoptera (Clarke et al. 2014; Marquina, Andersson, and Ronquist 2019). Thus, we also tested a primer pair targeting the 16S gene, which has been shown to have better taxonomic coverage for Hymenoptera and a range of other other insect orders (Diptera, Coleoptera, Lepidoptera, Hemiptera, Collembola)(Marquina, Andersson, and Ronquist 2019).

The final volume of PCR reaction was 20 μl, consisting of 2 μl DNA extract, 10 μl MyTaq HS Red Mix (Bioline Reagents), 1 μl forward primer (10 μM), 1 μl reverse primer (10 μM) and 6 μl MQ-H_2_O. PCR conditions are as follows: 96 °C for 7 min, followed by 40 cycles of 30 s at 96 °C, 30 s at 46 °C and 45 s at 72 °C, followed by 72°C for 5 min. The success of PCR reactions were evaluated by visualisation of the correct range of amplicon size on an electrophoretic gel running 2 μl of the PCR product. The PCR products were purified using columns MinElute PCR Purification kit (QIAGEN, Germantown, United States). In all PCR amplification steps a negative control was included (replacing the 2 μl of DNA by MQ-H_2_O) and checked on electrophoretic gel together with the amplified amplicons.

The purified PCR products were sent to the Swedish National Genomics Institute (NGI) in Uppsala for indexing, library preparation and sequencing. The protocol followed by NGI is described briefly as follows. Indexing was performed using the Adapterama indexing scheme (Glenn et al. 2019), using a 5 cycle amplification step, followed by bead (MagSI NGS plus beads) purification with a ratio of 1:1. Final libraries were eluted in 20 ul Elution Buffer. Libraries were quantified using a Quant-iT HS DNA assay and normalized by pooling to equimolar concentration (4nM for 16S samples and 10nM for COI samples). The 16S pool was subjected to an additional size selection using 0.7x beads to remove any traces of unspecific sizes at low molecular weight. The two prepools were combined at equimolar levels and the concentration of the final pool determined by qPCR using the Kapa library quantification kit from Roche. The pooled libraries were sequenced on Illumina platform (MiSeq, 2×300 PE) with a final addition of 10% Phi-X spike-ins.

### 2.4 Molecular species identification

Sequences were demultiplexed based on primers and the dual indices. Primers were trimmed with cutadapt (v. 3.4, Rognes et al. 2016) in paired-end mode, allowing for a maximum error rate (mismatches) of 10%. Sequences where the primers were not found or sequences shorter than 100 bp after trimming were discarded. Trimmed reads were processed with DADA2 (v. 1.18.0, Callahan et al. 2016). Based on the visual inspection of the quality profiles, the reverse reads were trimmed to 260 bp for COI and 240 bp for 16S, leaving a sequence overlap of 51 and 67 bp, respectively. The forward reads were not trimmed as the sequence quality stayed high over the full length. The trimming was followed by an error filtering, allowing for two expected errors in the forward reads and 4 expected errors in the reverse reads. We used the same settings for both barcodes. Finally the sequences were denoised and merged using the DADA2 algorithms. Chimera filter was performed using the *removeBimeraDenovo* function. Amplicon Sequence Variants were further clustered into OTUs at 99% identity using vsearch.

Taxonomic assignment of the COI sequences was done using the SINTAX (Edgar 2016) implementation of the RDP classifier (Wang et al. 2007) in vsearch. We used the MIDORI 2 database v. GB243 (Leray, Machida, and Knowlton, *in prep*) as the reference database, pre-formatted for SINTAX. For 16S, no reference database was available wherefore we created our own reference database. In short, we downloaded the full EMBL short read nt database (excluding organisms specific, synthetic, transgenic, unclassified and environmental sequences) and used ecoPCR from ObitTools3 (Boyer et al. 2016) to extract all sequences amplified by our 16S primers *in silico*. We discarded sequences that did not have a complete family, genus and species assignment and clustered the remaining sequences at 99% identity using vsearch. The taxonomic annotation of the sequences was harmonised against the gbif backbone using the rgbif package (Chamberlain et al. 2021) to resolve synonyms and avoid discrepancies caused by abbreviations added to species names such as *sp*. (species) or *cf*. (confer). We then checked the congruence of the taxonomic annotation in all non-singleton clusters. If incongruences occurred at the family level the cluster was excluded, if incongruences occurred at the genus or species level, the respective levels were set to *NA* & *sp*., respectively. Finally we formatted the reference database for SINTAX and assigned the taxonomy as for COI. The final 16S reference database contained 72.696 unique sequences.

Taxonomic assignments with probability of below 60% were set to NA. Subsequently, we removed all sequences with no taxonomic assignment at phylum level (40% of the sequences accounting for 14% of the reads for COI and 67% of the sequences accounting for 9.5% of the reads for 16S). We also discarded all human sequences and all sequences present in the blank. However, the blank sample contained only 11 sequences for COI and 9 Sequences for 16S.

### 2.5 Data analysis

For the comparison with the traditional surveying methods we only included sequences that retained a species level assignment after filtering (for eDNA) and only species that could be assigned to species level (for traditional monitoring). Species assignments from all data sources (16S, COI, and monitoring data) were harmonized against the gbif backbone taxonomy prior to species comparison. We checked the species for improbable detection and excluded three Lepidoptera species that are not occurring in Sweden nor in any neighbouring countries.

To assess the ability of the eDNA-based data to separate between sampling sites we calculate species turnover between the sampling sites for the COI data, we normalize the abundance matrix using the *varianceStabilizingTransformation* function from the DESeq2 R package (v. 1.30.1, Love, Huber, and Anders 2014). This transformation is an alternative to rarefaction (McMurdie and Holmes 2014). We calculated the bray-curtis dissimilarity based on the transformed matrix keeping all sequences and visualized the distance matrix with an NMDS plot. Statistical significance of the clustering was tested using the adonis2 function in the vegan package (Oksanen et al. 2020) with the formula *distance ∼ Station* (counting the three landscapes at the Pollinator site separately). We tested the clusters at the Pollinator site separately, including a habitat effect *(distance ∼ Station + habitat)*.

To estimate the sequence diversity at the different sampling locations, species accumulation curves for each site were estimated using the *specaccum* function using the ‘random’ method (providing the expected species number for n sites) and plotted against cumulative air volume by multiplying the number of samples by the volume of air collected for each sample at the different stations. We estimated species accumulation curves separately for each marker and including all sequences as well as only sequences assigned to the phylum “Arthropoda”.

If not stated otherwise, all data analysis, statistics and visualizations were performed in R v. 4.0.4 (R Core Team 2021) using these additional packages: *tidyverse* (Wickham et al. 2019), *eulerr* (Larsson 2020), *ggpubr* (Kassambara 2020), *phyloseq* (McMurdie and Holmes 2013) and *DECIPHER* (Wright 2016).

Upon publication, all scripts will be published on GitHub and all data published on Figshare. The raw sequencing data will be deposited on GenBank.

## 3 Results

### 3.1 Detected species

For COI, across all samples, 1277 sequences passed quality control and could be assigned at Phylum level or lower (with a minimum probability of 60%) while the 16S primers recovered 163 sequences that could be assigned at least to Phylum level. The number of recovered sequences differed greatly between the markers (Figure 2), with the differences primarily stemming from primer specificities: COI sequences were dominated by fungal taxa, with a total of 385 sequences identified as Basidomycea and 601 species as Ascomycetes. In total, the COI primers picked up sequences from 14 Phyla while the 16S primers almost exclusively amplified Arthropods (with a few sequences belonging to three other phyla). For Arthropods, the number of sequences was more balanced between both markers, with COI recovering 180 sequences assigned to “Arthropoda ‘‘ versus 152 sequences for 16S. The majority of recovered Arthropod sequences belonged to the class Insecta but sequences from six arthropod classes were detected across all samples and markers. For one of the classes, Hexanauplia, only one sequence was detected which was assigned to the order Cyclopoida, a subclass of Copepods (100% match on BLAST to an environmental Cyclopoida sample). However, the sequence also perfectly matches the sequence of *Fujientomon dicestum* belonging to the order Protura (Coneheads) and class Entognatha - a group of small soil dwelling arthropods. It is therefore unclear which class was detected. From the insect class 117 sequences were detected with the COI primers versus 130 sequences from 16S. The sequences represented 12 insect orders with the majority of sequences belonging to the four largest orders: Diptera (Flies), Lepidoptera (Butterflies and Moths), Coleoptera (Beetles) and Hymenoptera (Bees, Wasps and Ants).

**Figure 2.**
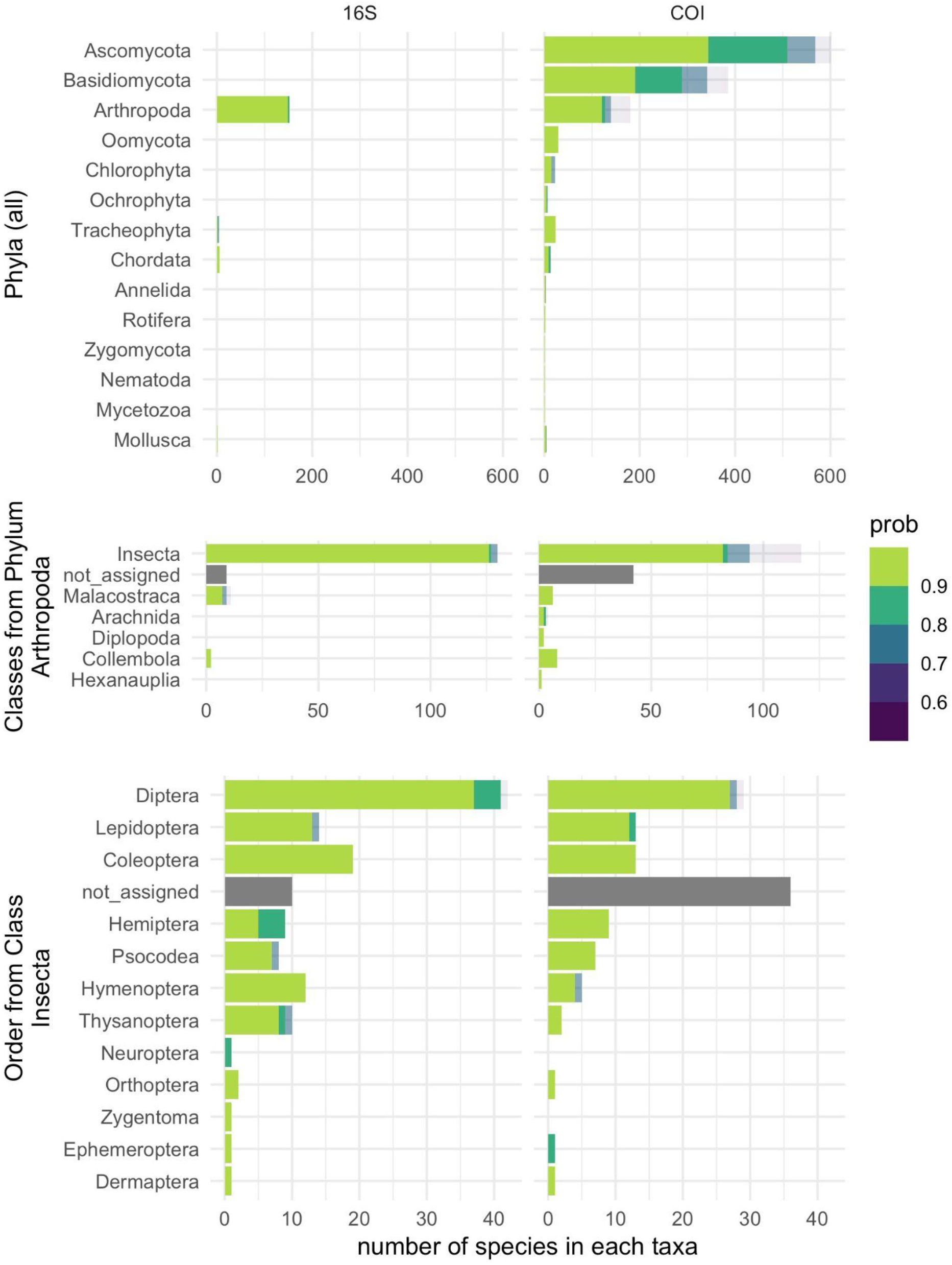
The distribution of taxa detected from airborne eDNA, for both the COI and the 16S markers. Only sequences that were assigned a taxa with at least 60% probability at this taxonomic level were considered and the number of sequences in each probability class are colour coded (probabilities are binned to the nearest ten percent). ‘not_assigned’ shows the number of sequences that could not be assigned to a taxa at this taxonomic level (probability < 60%). The first row shows the number of sequences in each phylum, the second row shows the number of sequences within the phylum Arthropoda (by class) and the third row shows the number of sequences in the class Insecta (by order).

While not the target group of this study, both primers also recovered sequences from vertebrates, detecting nine species belonging to eight orders: a pool frog, three bird species (the common wood pigeon, a fieldfare and domestic fowl) and five mammals (cattle, dog, European hedgehog, red squirrel, short-tailed field vole). We also detected the European rabbit (*Oryctolagus cuniculus*) but this sample was excluded from the analysis as it wasn’t taken during one of the regular sampling schemes. All detected vertebrate species are common in the region and three are domestic species (cattle, domestic fowl and dog). Note that the chosen markers cannot distinguish dog from grey wolf but the latter is very rare in the region and therefore highly unlikely.

### 3.2 Comparison to traditional sampling

Figure 3 and Figure 4 show the comparison between species detected from eDNA and with traditional sampling methods (light trapping and transect surveys). Only sequences belonging to the kingdom Animalia were included, and only those that could be identified to species level (60% probability of the assignment). In total 50 unique species of insects could be assigned a taxonomy down to species level for COI and 43 species for 16S. Across both markers, 85 unique species of insects were detected. Across the ten samples taken at ten consecutive weekdays at the Moth site, seven species of moths were detected with COI and two species with 16S. During the same period, 48 moth species were sampled in the light trap. Five species were uniquely detected from eDNA while four species were detected by both eDNA and light trapping.

**Figure 3.**
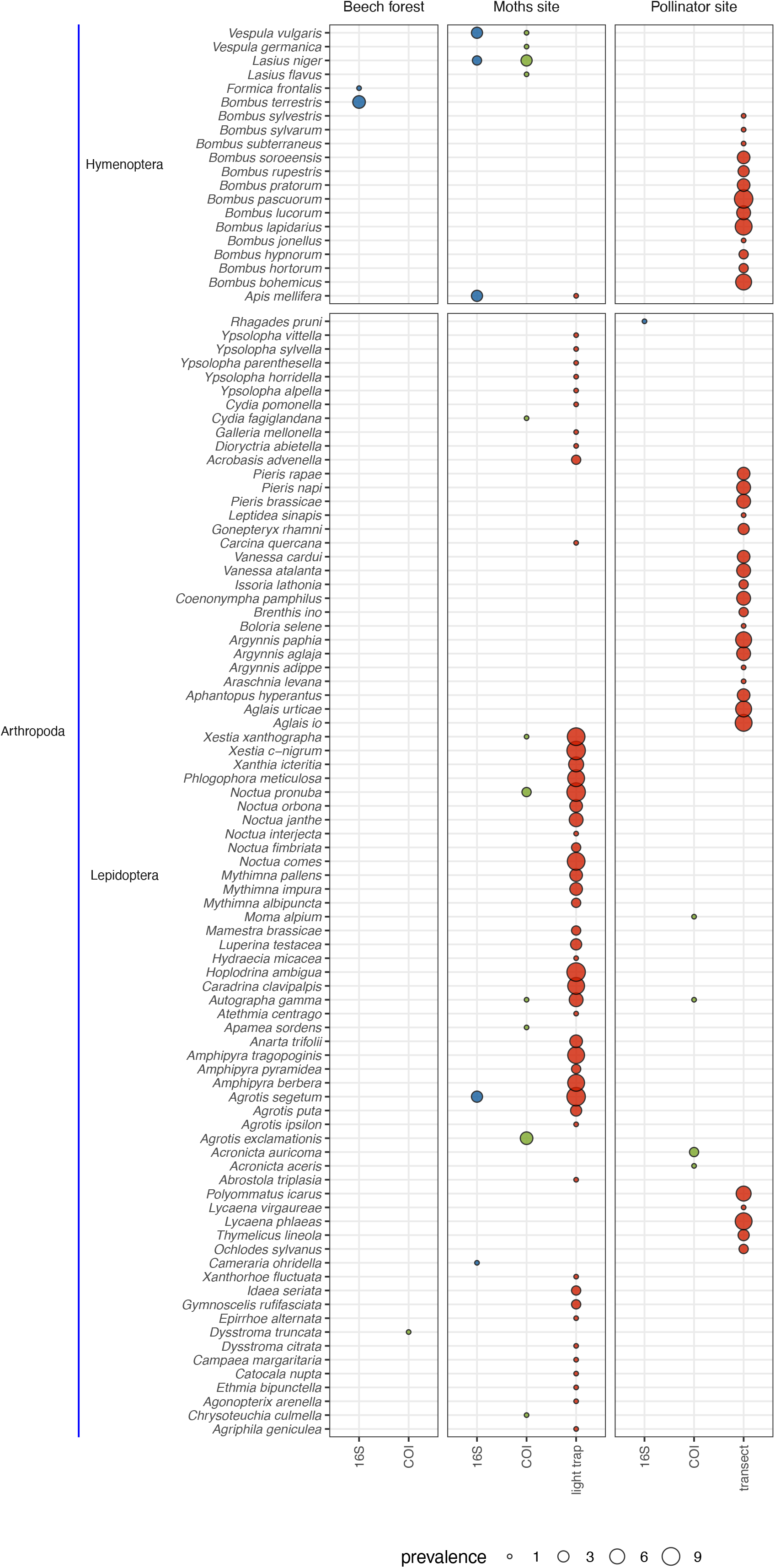
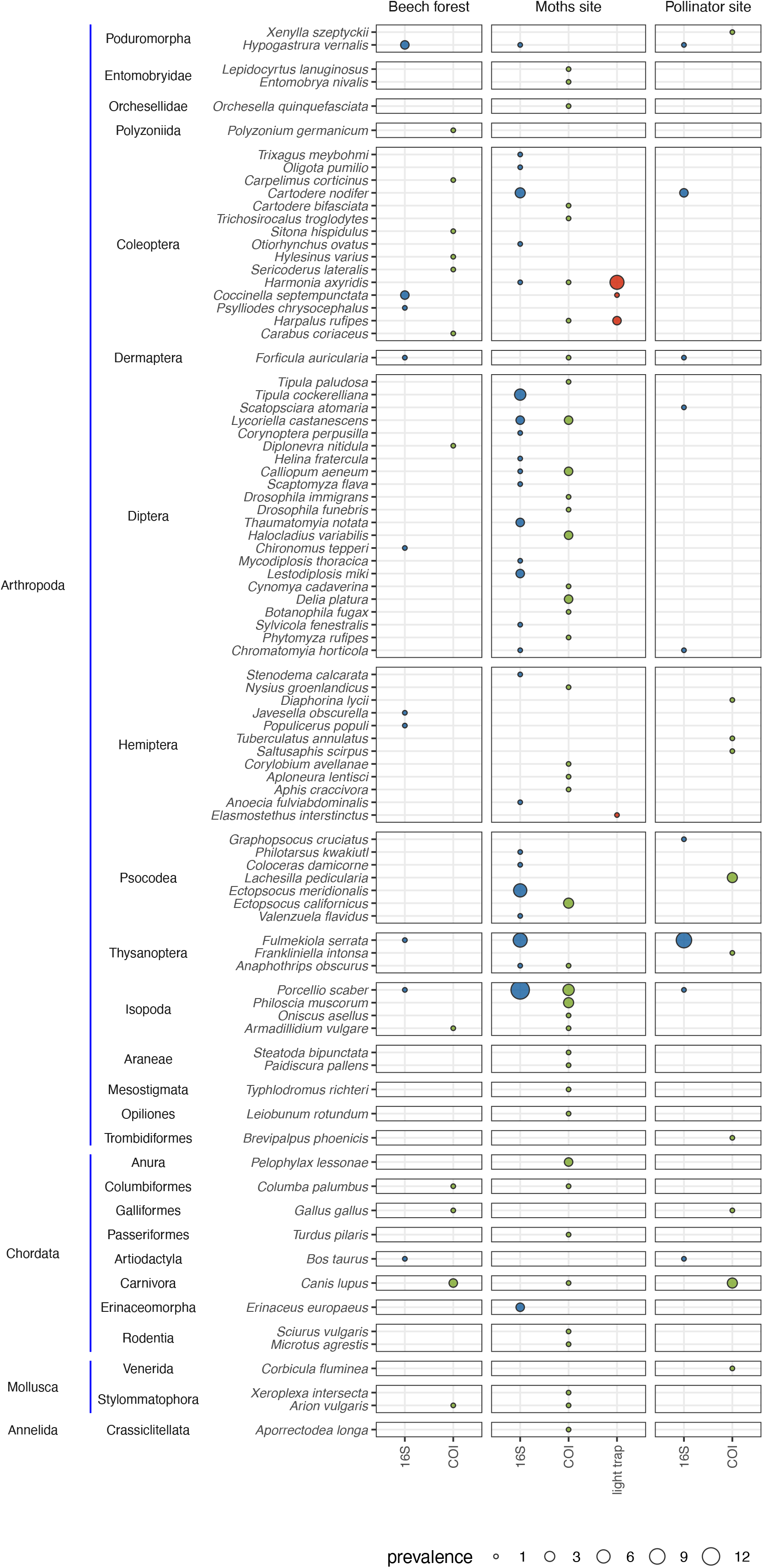
Bubble chart showing which species were detected in what locations, both with eDNA (separately for each marker) and in the surveys. Only Species belonging to the animal kingdom are shown. The size of the bubble is proportional to the prevalence of the species across samples at that location (maximum 5 at Bokskogen, 10 at the Moth trapping site and 15 at the pollinator surveying site). Species are ordered phylogenetically and the ranks phylum, order and species are given.

**Figure 4.**
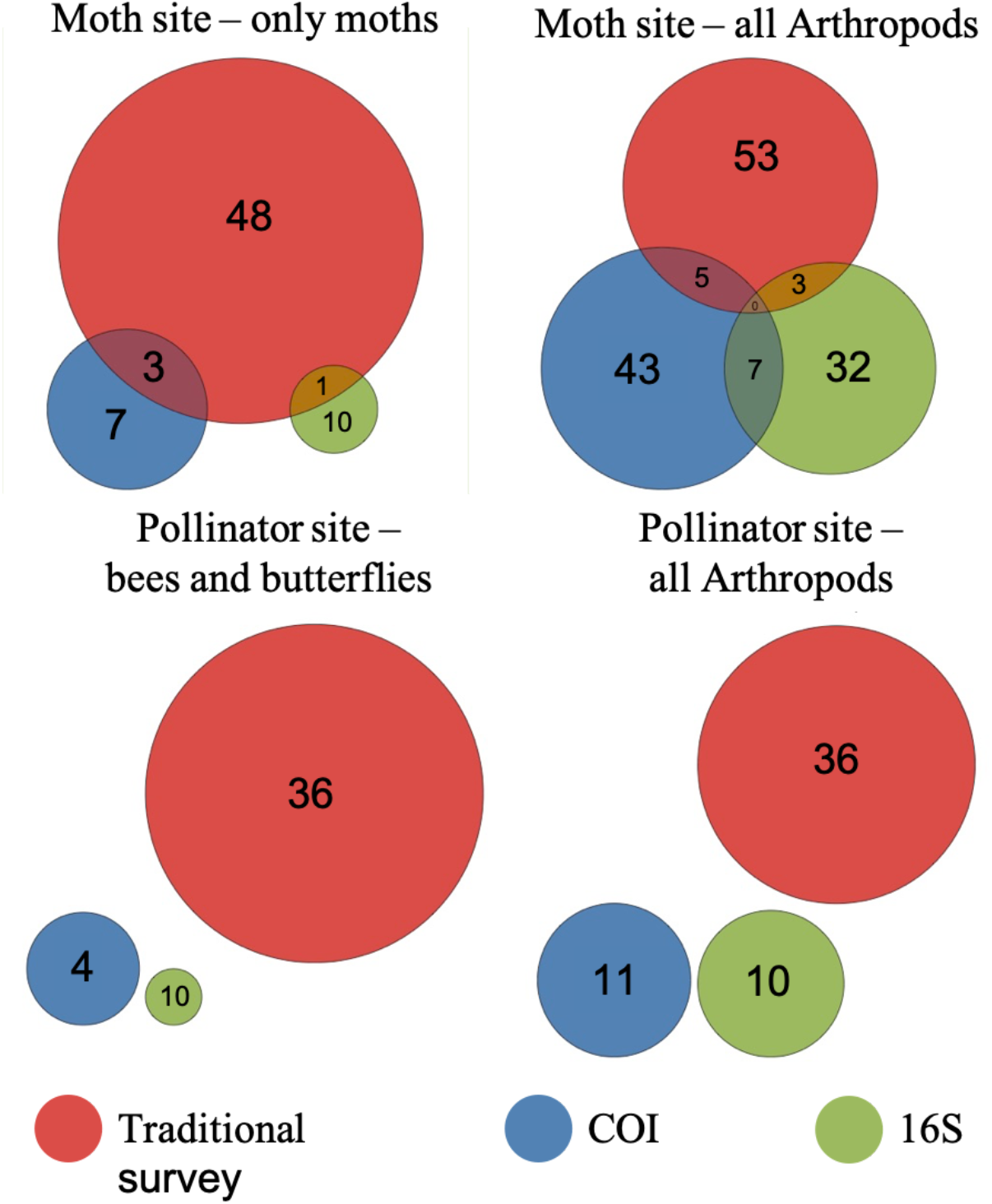
Euler diagram presenting the same data as figure 3 but highlighting the number of shared and unique species that were detected for both markers and the traditional surveying methods, respectively. The left panel for each surveying scheme counts only species that were targeted by this monitoring scheme (Lepidoptera for Moth and Lepidoptera and Apidae for the Pollinator survey), while the right panel counts all Arthropods. During the light trapping non-moths arthropods were also recorded if encountered in the trap. The pollinator survey did only record butterflies and bees.

At the Pollinator site, butterflies and wild bees were surveyed using transect counts. When comparing only the taxa targeted by the transect counts, the order Lepidoptera and the family Apidae (Hymenoptera), five species were detected by eDNA (four from COI and one from 16S) versus 36 species from the transect counts. No common species were detected.

When we expand the focus on all Arthropods the picture changes: At the moth site, 67 species of Arthropods were detected by eDNA (43 for COI, 31 for 16S, 7 in common) while 53 species were detected in the light trap. The latter includes five non-moth arthropods which were also recorded in the light trap. At the Pollinator site a total of 20 arthropod species were detected with eDNA versus 36 species in the transect counts. During the transect counts no other arthropods were recorded.

### 3.3 Ecological information

We calculated the Bray-Curtis dissimilarity between all samples across the three sampling sites based on all sequences that passed the quality control, for COI. The NMDS showed a clear grouping of the samples by sampling location (Figure 5) and a Permutational Multivariate Analysis of Variance test showed that the clustering was significant (*distance ∼ Station*, F_4,29_ = 3.08, R2 = 0.33, p = 0.001). At the Pollinator site, the samples were taken in clusters of five, in three different landscapes which were between 20 - 50 kilometers apart (SF - SI: 20km, SF - SB: 50 km, SB - SI: 40 km). Here, too, the samples from the same landscapes formed distinct clusters and the clustering was significant (*distance ∼ Station + Habitat*, F_2,10_ = 1.7, R2 = 0.22, p = 0.001). However, the different habitats could not be told apart by their species composition (p = 0.24). For 16S the data were too sparse to calculate a meaningful dissimilarity.

**Figure 5.**
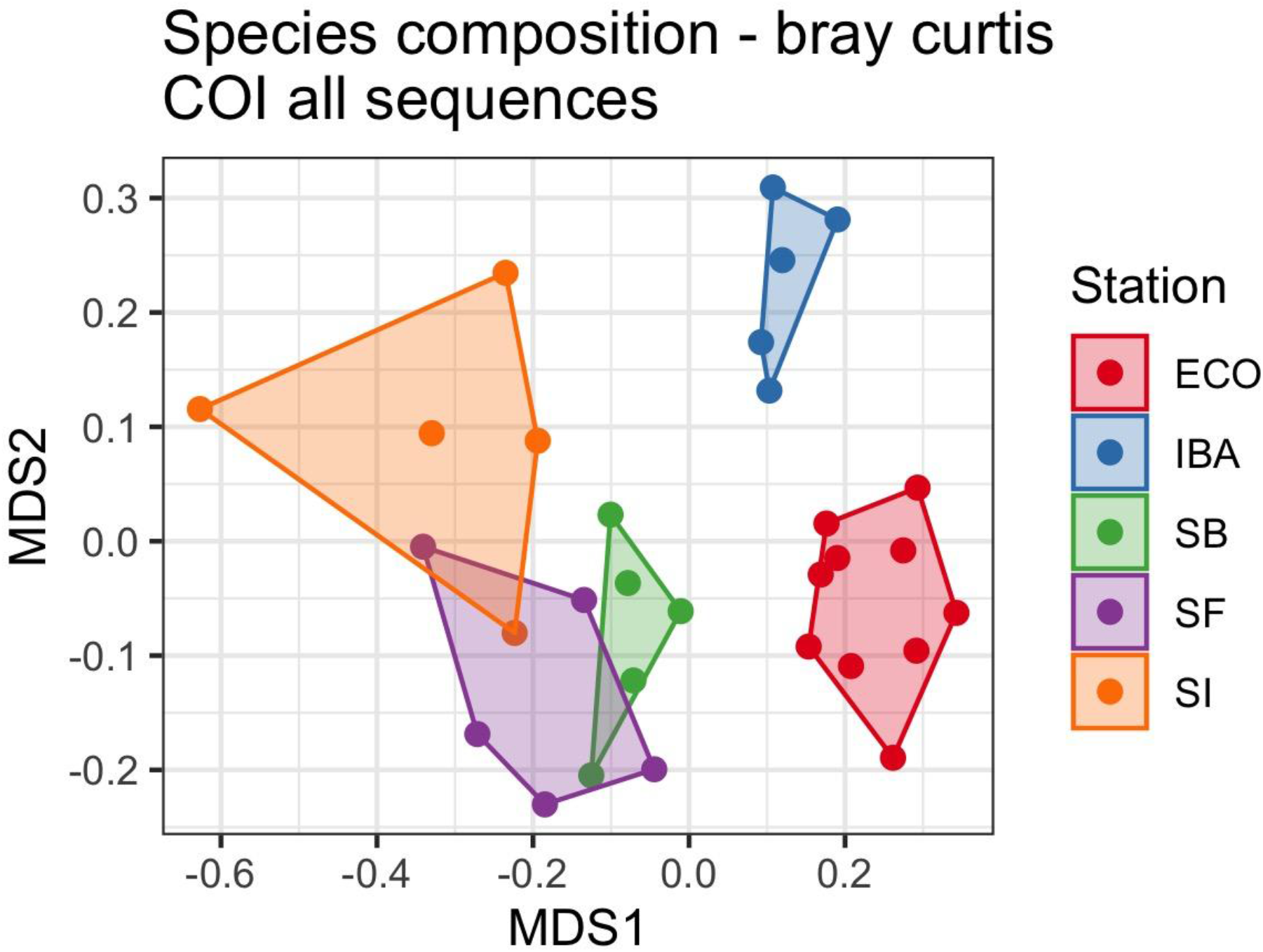
NMDS plot of all samples, coloured by sampling site. The three locations at the Pollinator site are coloured separately. The NMDS is based on bray curtis dissimilarity. Prior to calculating the distance, counts were normalized using the *varianceStabilizingTransformation* function from the DESeq2 R package.

We also calculated the cumulative sequence richness at each sampling site (Figure 6). The Moth site had remarkably higher diversity than the Pollinator site, both for COI and for 16S and both if all sequences were considered or only those identified as Arthropods. The Beech forest showed a similar trend than the Moth site but due to the lower total sampling volume (30 m^3^ versus 90 m^3^ of air) the pattern for the Beech forest is hard to assess.

**Figure 6.**
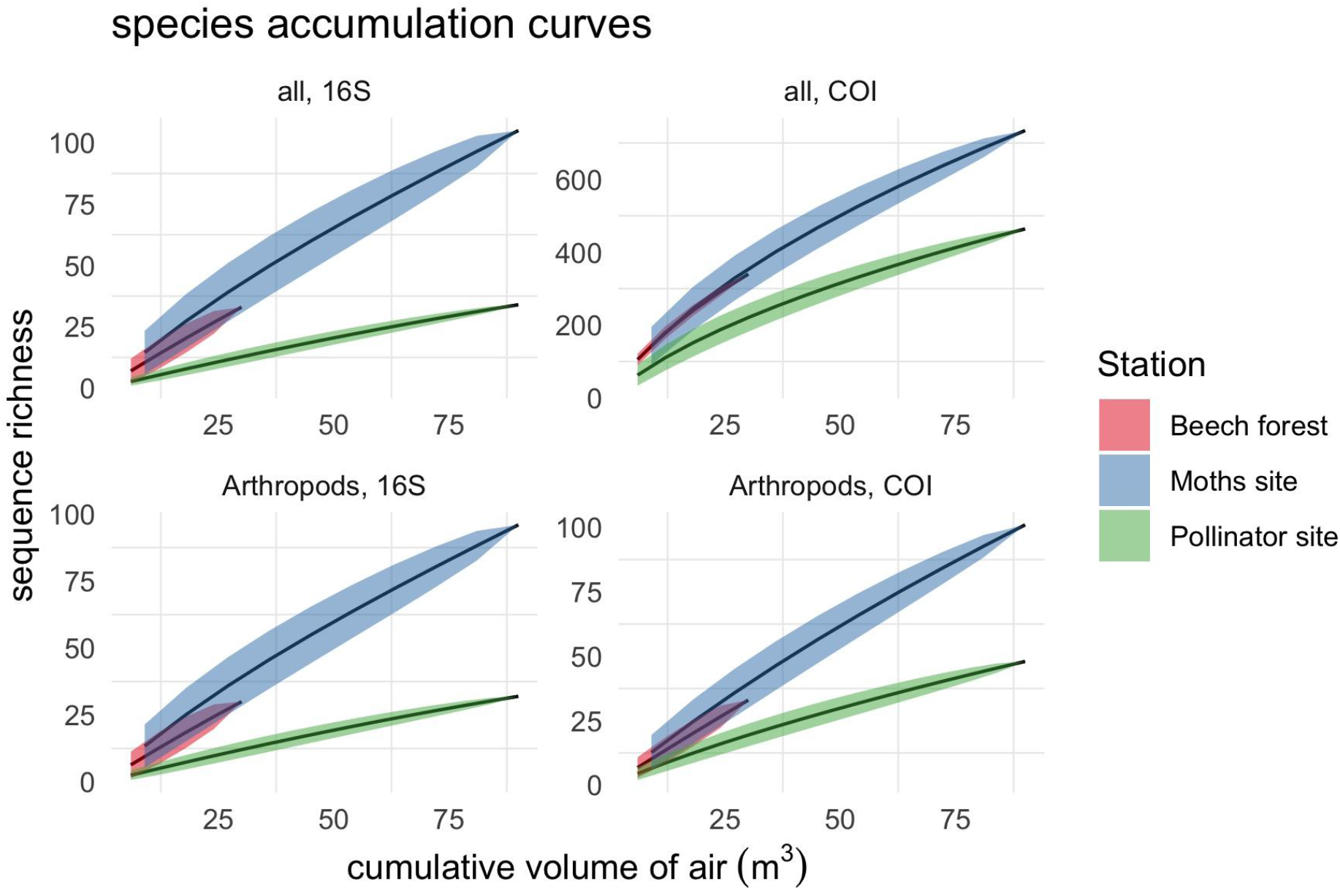
Sequence accumulation curves for the three sampling sites. The accumulation curves show the expected (average) number of species detected for a certain volume of air, wenn samples are added at random. We show the accumulation curves separately for both marker (left column - 16S, right column COI) and for all sequences (upper row) and only sequences identified as arthropods (lower row). The Moth site and Pollinator site have the same total volume of air as the Moth site had 10 samples but each with 9 m^3^ of air while the Pollinator site had 15 samples but only 6 m^3^ per sample. Note the different scales on the y-axis for each panel. Shaded ribbons represent one standard deviation around the mean.

## 4 Discussion

This study represents a proof-of-concept that insects, and many other species groups can be detected from airborne environmental DNA.

The comparison with the traditional monitoring data has shown that - as of now - airborne eDNA fails to detect many species, especially butterflies and wild bees (with some more success for moths).

Considering all Arthropod groups though, a substantial diversity of species was detected (101 unique Arthropod species could be identified to species level), showcasing that eDNA from insects and other arthropods is present in the air and can be targeted for species detection and monitoring. Moreover, the unspecific amplification of the COI primers showed that eDNA from a wide variety of taxa is present in the air, including vertebrates, molluscs, annelids, plants, green algae, oomycetes and many species of fungi - both ascomycetes and basidiomycetes (the former representing many filamentous and single celled species while the latter includes most species commonly known as ‘mushrooms’ that produce airborne spores). This suggests that airborne eDNA could be a powerful tool for a comprehensive biodiversity assessment of terrestrial ecosystems.

### 4.1 Metabarcoding primers for airborne eDNA

Metabarcoding of insects is challenging as insects represent a mega-diverse order and designing primers that target all insect species simultaneously is difficult. In this study we chose two primer pairs that were both developed for the metabarcoding of insects and have been found to have little taxonomic bias and broad taxonomic coverage (Elbrecht et al. 2019; Marquina, Andersson, and Ronquist 2019). However, the primers have been developed for insect bulk samples (from Malaise traps) where co-amplification of non-target groups is not a major concern. In contrast, the eDNA samples collected in this study were likely dominated by non-target DNA (fungi, bacteria and other microorganisms - as it is typical for eDNA from other sources as well (Turner et al. 2014) and co-amplification was a substantial problem. The sequences amplified by the COI primers were dominated by fungi (75% of all reads) and microscopic Fungi and spores represent living organisms, many composed of thousands of living cells. As such, they can easily dominate the pool of mitochondrial DNA sequences. This leads to a significant reduction in sequencing power available to detect low-abundant extra organismal DNA from insects. The 16S Primers suffered only low levels of co-amplification of non-target taxonomic groups, amplifying almost exclusively 16S sequences from Arthropods. Yet, the primers seemed to amplify a large number of seemingly random sequences that did not appear to be 16S sequences and could not be assigned any taxonomic annotation, even at phylum level. The sequences also did not cluster together phylogenetically, producing a tree with many long branches and little structure (Suppl. Figure 1). We therefore suspect that the primers might amplify random genomic regions and sequences from this accounted for 10% of all reads, again, entailing a loss in sequencing power. While these problems likely limited the sensitivity of our study, it entails that optimizing primers and/or choosing primers targeting specific insect groups has a large potential to increase the sensitivity of airborne eDNA sequencing in the future.

### 4.2 Aerosol sampling

A range of techniques are available to collect aerosols (Dybwad, Skogan, and Blatny 2014; Haig et al. 2016) including eDNA carrying particles. The samplers differ in whether they sample particles actively (by sucking in air) or passively (sedimentation plates). Passive air samplers have been used successfully in two recent pilot studies targeting airborne eDNA from honey mesquite (a non-anemophilous plant) (Johnson, Cox, and Barnes 2019; Johnson et al. 2021). They are less costly and do not rely on a power source, and can therefore be deployed for extended amounts of time. However, quantification and standardization of the samples are uncertain, they oversample large aerosol particles and have generally poor collection efficiency. Active air samplers have higher collection efficiency, a more uniform and representative sampling of aerosol particles and a possibility to control sample volume, by controlling the flow rate and time of sampling. This allows a standardized sampling, and the sampling of large volumes of air in short time spans, both of which is likely to be advantageous for future applications of airborne eDNA sampling.

In this study we used the Coriolis µ for the collection of eDNA carrying aerosols, which samples into liquid (facilitating molecular downstream analysis), has a high flow rate (up to 300 L*min^-1^), is portable, easy to decontaminate and can be battery driven. We ran the sampler at the highest flow-rate for 10 min, which was the maximum duration without a long-sampling option that replaces evaporated liquid during the sampling. To collect larger samples, we pooled two - three samples, reaching a collection volume of six to nine m^3^ of air / sample. This represents a cube of a little more than 2 × 2 x 2 meters, which might not be enough to collect sparsely occurring eDNA carrying particles. In line with recent findings that show that terrestrial vertebrate detection from streams can be substantially improved through larger and replicated sampling (Leese et al. 2020), replicated larger samples of air might increase species detection substantially.

A range of alternative aerosol samplers exists, such as the BioSampler (that can be run for multiple hours without evaporation), the Kärcher impinger (Šantl-Temkiv et al. 2017 - with a flow rate of 3000 L min), the Anderson liquid stage impactor (that allows for size fractionated sampling) and many more. All have their advantages and disadvantages and no standard exists that is suitable for all types of biological aerosol collection. Which sampling strategy is most efficient to retrieve eDNA from the air will need to be examined.

### 4.4 Origin and fate of airborne eDNA

One important question for the species detection and monitoring of DNA is the origin of the signal (Barnes and Turner 2016): If extra-organismal DNA is detected, it could have travelled long distances before being captured weakening the link between detection and presence of a species at the sampling site (Harrison, Sunday, and Rogers 2019). While this is true for eDNA in water, too, where it can be transported with currents and through river networks (Deiner et al. 2016), many aquatic ecosystems are relatively confined (ponds, lakes, catchment areas - with the ocean being the exception). In contrast, the atmosphere represents a circumglobal ‘ocean’ of air, with high-velocity currents (winds) and strong turbulence. Depending on their size (or aerodynamic diameter), aerosols particles can stay airborne for extended periods of time and potentially be transported over long distances (Emerson et al. 2020). As an example, the HYSPLIT model (Draxler and Rolph 2010) can be used to calculate the trajectories of the air masses that reached our sampling location at a certain time and height. Figure S2 shows the calculated trajectory for the air masses that reached the Ecology building in Lund (Moth site) at a height of 10 m on August 29^th^ at 8 a.m. The 48h backward trajectory projects an origin from the air masses reaching from Wales in the West to near the city of Frankfurt in Germany to the South, to the western border of Ukraine to the East. Small particles could easily stay airborne for 48h putting a large uncertainty on the origin of the collected DNA. Yet, our results show a distinct species composition for the three sampling locations.

Samples collected at the same station strongly clustered together - and even the three subclusters at the Pollinator site which were separated by only 20 - 50 km, showed a strong grouping.

Contrary to what one could expect, this suggests that airborne eDNA carries a strong local signal. This could be explained by (1) dilution (concentration of eDNA will decrease rapidly with distance from the source (Harrison, Sunday, and Rogers 2019), and low viscosity and high turbulence of air might exacerbate this effect) (2) fast degradation (UV degrades DNA rapidly (Lindahl 1993) and very efficiently, and especially small particles might be exposed to strong UV during daylight) or (3) low transportability of the particles (if the bulk of the signal comes from larger particles that stay airborne only for short periods at the time). In our study we speculate that for COI, the local signal was driven by fungi (spores and microscopic) and other living microorganisms (microalgae, Oomyctes), which accounted for the bulk of the sequences. Fungal spores are large (for aerosols) and a local to regional signal from fungal spores has been previously found (Abrego et al. 2018; Olsen et al. 2019). We couldn’t assess the clustering of the sequences that originated from extra-organismal DNA alone (Animal Kingdom) as the matrices were too sparse to calculate meaningful dissimilarities. The locality of the signal from extra-organismal airborne eDNA thus remains to be studied and might well differ between organisms, aerosol size fractions and environmental conditions.

An interesting observation is also that the diversity of captured sequences differed markedly between sampling locations. At the Moth site, two to three times more sequences were detected for the same volume of sampled air than at the Pollinator site (Figure 6), for both markers and regardless if all sequences or only Arthropod sequences where considered. The Beech forest followed the pattern of the Moth site but as the total sampled volume of air was only one third of the other sites, the pattern is less clear. Given the design of the study we can only speculate why these differences occurred but there are some interesting potential explanations. First, the Moth site and the Beech forest were both situated in relative proximity to each other (∼20 km) and about 200 km south of the Pollinator site. The Moth site is located in a city, and the Beech forest is in large parts a protected area rich in species, while the Pollinator site(s) are agricultural sites embedded in a matrix of species poor production forest (mainly spruce monocultures). Urban forests and protected forests have been found to be more diverse than production forests (Korhonen et al. 2020) although a recent study from Finland found that fungal diversity in the air declined steeply with urbanization (Abrego et al. 2020).

Putting the Beech forest aside (where the pattern is difficult to assess), other explanations could also play a role: The sampling at the Moths side was conducted in the mornings, and the weather during the sampling period was variable (with some calmer and some windier days, and quite a few days with cloud cover), while the weather conditions during the sampling at the Pollinator site were windstill and sunny. This could suggest that either UV degradation plays an important role and airborne eDNA should be sampled on cloudy days or before sunrise or that windy conditions increase the diversity of eDNA in the air. The latter could be the case if much of the eDNA sampled from the air was caused by resuspension of top-soil - which might be a likely source and could explain the detection of soil organism such as the detected earthworm (*Aporrectodea longa*), spanish slug (*Arion vulgaris*) and land snail (*Xeroplexa intersecta*). Finally it is also possible that the sampling height has a significant influence on the capture bioaerosols, as the wind speed at ground level is zero (due to surface friction) and increases with increasing height above the ground (wind speed gradient). As a result, higher sampling locations might contain a more diverse mixture of aerosols from a larger region.

### 4.5 Potential applications

In the wake of a global biodiversity crisis, the need for more, more accurate and faster biodiversity information becomes ever more apparent. The Convention on Biological Diversity has been ratified by 192 Nations, the United Nations sustainable development goals call to “Protect, restore and promote sustainable use of terrestrial ecosystems […] and halt biodiversity loss” (Goal 15), and the European Union’s Biodiversity strategy for 2030 “aims to put Europe’s biodiversity on the path to recovery by 2030”. To achieve these goals, new methods to monitor the world’s biodiversity are indispensable, especially for the vast majority of taxa that are not in the focus of traditional biodiversity monitoring. While only an imperfect first step, this study provides a proof-of-concept that Insects - the most diverse animal taxa on earth - leave detectable traces of DNA behind in the air. Moreover, we find that the same is true for a number of animal and non-animal phyla, including a range of vertebrates. This shows that species detection from airborne eDNA is possible - a result that could be the key that unlocks the access to a comprehensive and scalable method of terrestrial biodiversity monitoring at global scale.

The possible applications go beyond biodiversity monitoring alone: Insects can be devastating agricultural and silvicultural pests and the cost of invasive insects has been estimated to a minimum of US$ 70 billion per year globally (Bradshaw et al. 2016). Species-specific detection of insect pests from airborne eDNA could serve as early warning systems, support precision agriculture and be used to monitor goods at sensible points of entries (e.g. harbours, airports, logistic centres or shipping containers). Similar efforts are already underway in Sweden where suction-trap catches of aphids can help predicting aphid outbreaks (Jonsson and Sigvald 2016) and in Australia where the iMapPESTS project is developing mobile suction traps that monitor fungal spores and insects with molecular methods (https://ausveg.com.au/biosecurity-agrichemical/biosecurity/imappests/). Airborne eDNA sampling could allow for a simultaneous surveillance of multiple agricultural pests, including fungi, water molds and insects - three groups that all were detected in this study. The targeted detection of insects could also be applied to the monitoring and early detection of disease vectors, which encompass both, vectors of human pathogens (e.g. Malaria) and vectors of plant pathogens.

### 4.6 Future perspectives

While it is now clear that many species leave detectable traces of DNA behind in the air, future research is needed to realize the full potential of airborne eDNA bases methods. Critical next steps will be to optimize sampling and laboratory protocols to uncover the full breadth of diversity present in airborne eDNA, learning from the great advances that have come out of aquatic eDNA research over the last decade. We will also need to characterize the particles that carry eDNA in the air and to gain an understanding of the mechanisms governing the generation, transportation and degradation of these particles. Bioaerosol scientists have studied bioaerosols for decades and a vast body of knowledge around bioaerosols exists - giving airborne eDNA science a running start and an opportunity for fruitful new collaborations. The journey has just begun and many challenges will have to be overcome. But the many impactful applications make airborne eDNA an exciting new area of research and promising tool in our fight to monitor, understand and finally protect the biodiversity around us.

## Acknowledgment

We thank Prof Adam Kristerson for inspiring discussion at the start of this project and Dr. Celine Mericer for help with the OBITools platform. We acknowledge support from the National Genomics Infrastructure in Stockholm funded by Science for Life Laboratory, the Knut and Alice Wallenberg Foundation and the Swedish Research Council, and SNIC/Uppsala Multidisciplinary Center for Advanced Computational Science for assistance with massively parallel sequencing and access to the UPPMAX computational infrastructure.

## Supplementary Figures

**Figure S1.**
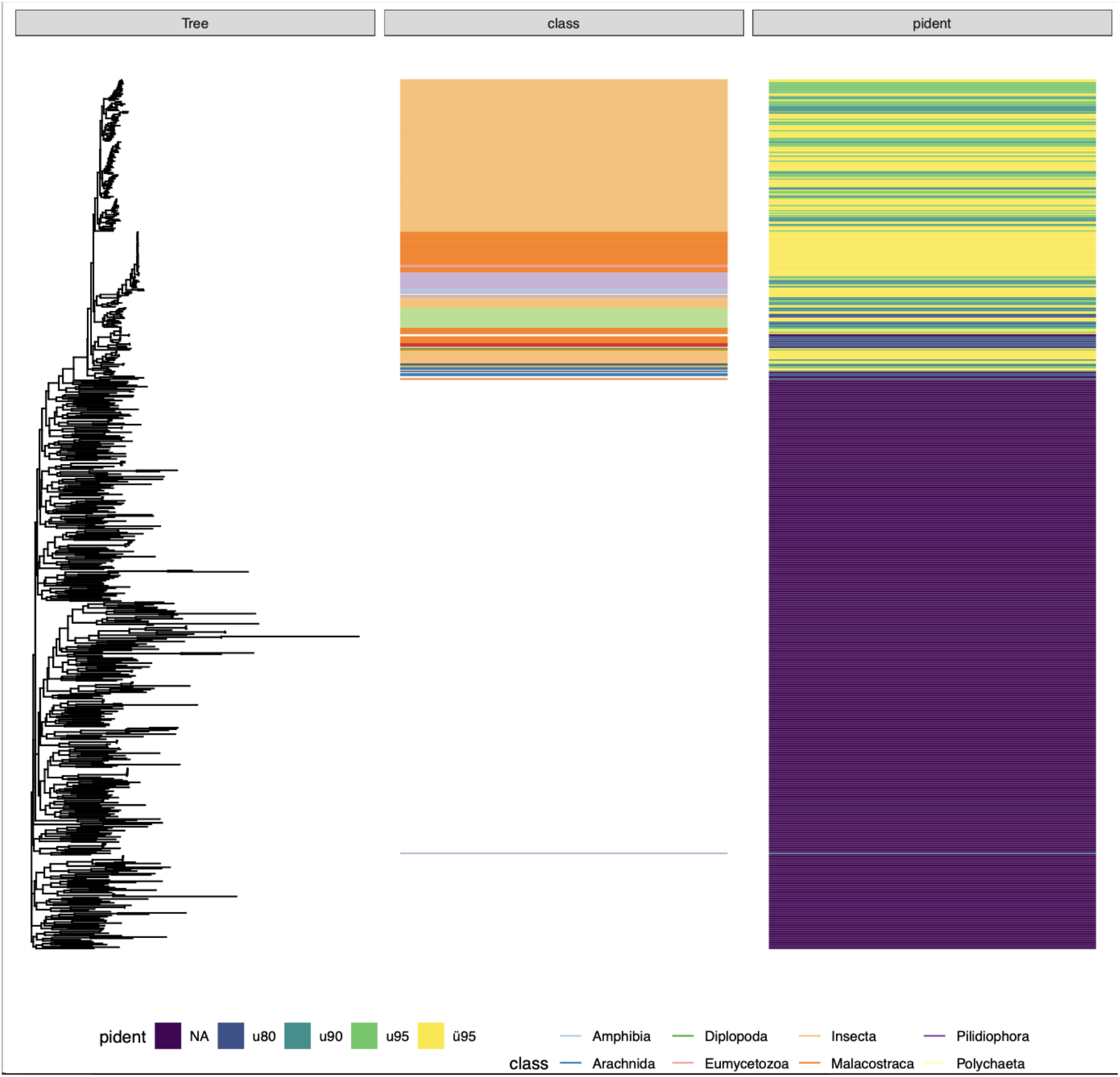
A neighbor-joining phylogenetic tree from all 16S sequences. Sequences were aligned with the Decipher R package and the tree was constructed using FatsTree with default values. The colours in the second panel show the taxonomic assignment at class level from the best hit and the third panel shows the percent identity of that hit. The tree shows two distinct clusters, an upper cluster with short branches and credible taxonomic assignment, and a lower cluster with very long branches and no taxonomic assignment at all. This suggests that the 16S primers could co-amplify random genetic regions but from what organisms is not clear.

**Figure S2.**
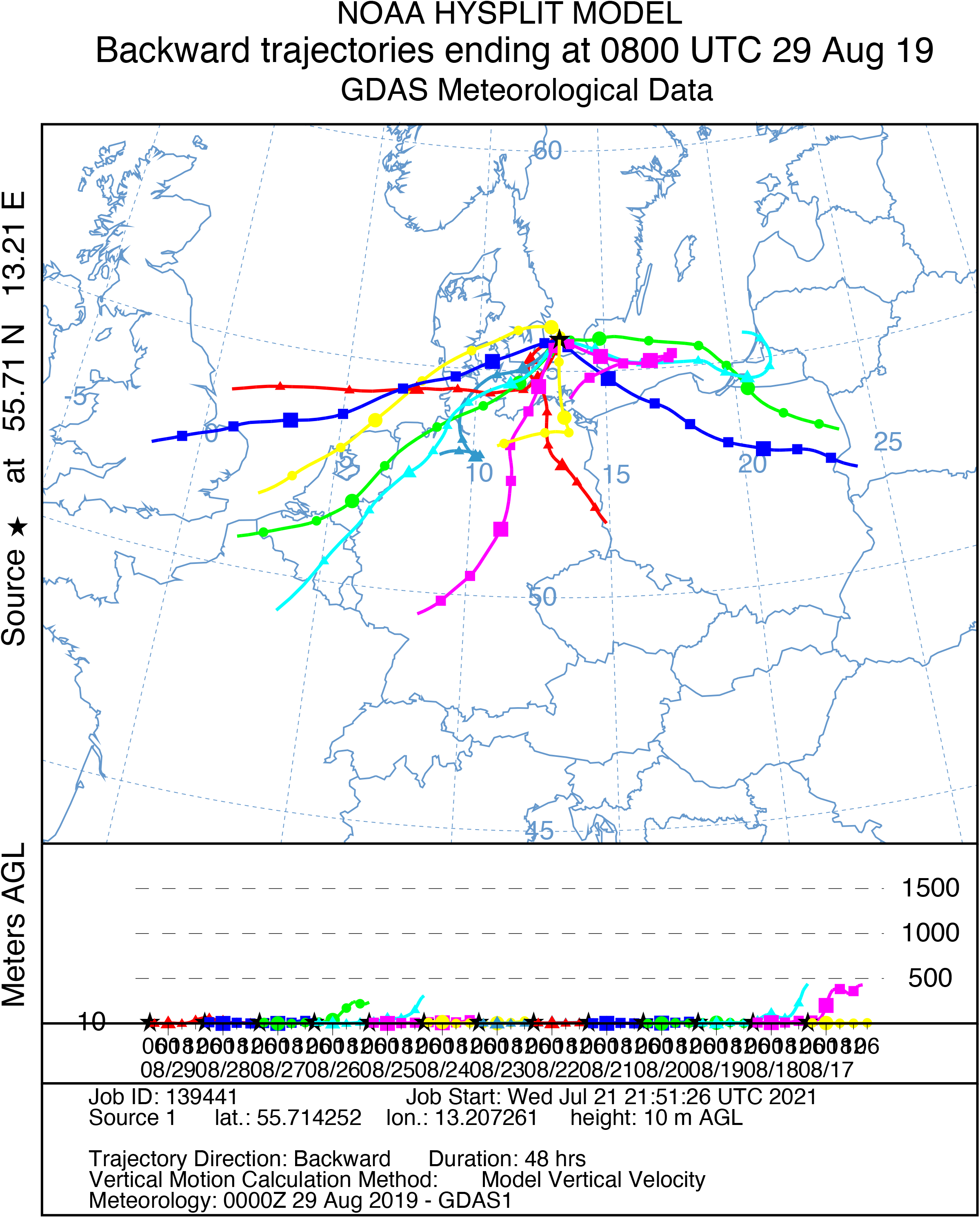
Backtraced trajectories from the air masses (48h) that reached the sampling location in Lund (Moth site) on the 29th of August 2019 (one of the mornings during which eDNA samples were collected). The trajectories were calculated using the HYSPLIT model [REF] for a sampling height of 10 m. The plot shows that the air reaching the sampling location at the time of sampling had diverse geographic origins and has traveled hundreds of kilometers in the previous two days.

## Notes

### Competing Interest Statement

The authors have declared no competing interest.

